# Nuclear bundle/cable containing actin during yeast meiosis

**DOI:** 10.1101/778100

**Authors:** Tomoko Takagi, Masako Osumi, Akira Shinohara

## Abstract

Actin polymerizes to form filaments/cables for motility, transport, and structural framework in a cell. Recent studies show that actin polymers are present not only in cytoplasm, but also in nuclei of vertebrate cells, and their formation is induced in response to stress. Here, by electron microscopic observation with rapid freezing and high-pressure freezing, we found a unique bundled structure containing actin in nuclei of budding yeast cells undergoing meiosis. The nuclear bundle/cable during meiosis consists of multiple filaments with a rectangular lattice arrangement often showing “feather-like” appearance. The bundle is immuno-labeled with anti-actin antibody and sensitive to an actin-depolymerizing drug. Like cytoplasmic bundles, nuclear bundles with actin are rarely seen in pre-meiotic cells and spores, and are induced during meiotic prophase-I. The formation of the nuclear bundles/cables is independent of meiotic DNA double-stranded breaks. We speculate that nuclear bundles/cables containing actin play a role in nuclear events during meiotic prophase I.

## Introduction

In the cytoplasm, as a cytoskeletal protein, actin polymerizes to form a filament (F-actin) for various cellular functions such as motility, division, phagocytosis, endocytosis, and membrane trafficking ^1^. Dynamics of cytoplasmic actin filaments are highly regulated by various factors in different environments. Actin is also present in nuclei ^2^. Actin monomer functions as a component of several chromatin-remodeling complexes for transcription and other nuclear events ^3–6^.

Recent studies showed that polymerized forms of actin are present in nuclei of various types of vertebrate and invertebrate cells ^7–12^. About forty-years ago, Fukui and his colleagues identified actin bundles in *Dictyostelium* and HeLa cells upon the treatment with dimethyl sulfoxide ^13–15^. In *Xenopus* oocytes, nuclear actin forms a mesh of filaments, which is involved in the protection of nucleoli from gravity-induced aggregation ^16^. In starfish oocytes, actin filaments promote the breakdown of nuclear envelope and, by forming a mesh, the capture of chromosomes by spindles in cell division ^17^. In mouse oocytes, actin filaments promote chromosome segregation during meiosis I and II ^8^. Somatic mammalian cells induce the formation of actin polymers transiently in a nucleus in response to stress; serum starvation, heat shock, and DNA damage such as DNA doublestrand breaks (DSBs). For serum starvation, the F-actin is involved in transcription by helping the activity of a transcriptional cofactor, MRTF (myocardin-related transcription factor) ^18, 19^. Nuclear F-actin also promotes repair of DSBs in mammalian and fruit fly cells ^7, 10,20^.

In budding and fission yeasts, actin is present in the cytoplasm as a polymerized form such as rings, patches and cables ^21–23^ as well as the filasome, which is a less-defined cytoplasmic amorphous structure containing F-actin ^24^. In budding yeast *Saccharomyces cerevisiae*, actin cables in cytoplasm play a role in the transport in mitotic budded cells. Previous electron microscopic (EM) analysis of cytoplasm in fixed mitotic cells revealed a linear actin bundle/cable containing multiple actin filaments ^25^. Actin polymers are also visualized by using a green fluorescent protein (GFP) fused with an actin-binding protein, Abp140, or by staining with an actin-specific peptide with a fluorescent dye such as phalloidin ^26^. These staining confirmed the presence of filaments/cables and patches containing actin polymers in cytoplasm of the yeast.

Actin polymers/cables are present also in of the cytoplasm of meiotic yeast cells ^27, 28^. The actin cables are induced during meiotic prophase-I (meiotic G2 phase) and form a network that surrounds the nucleus. The cytoplasmic actin cables attached to protein ensembles in nuclear envelope (NE) drives the motion of telomeres, thus chromosomes ^27, 29^. Although these cables are actin polymers, fine structures of actin cables are less defined. Previous EM observation of a chemically fixed cells undergoing meiosis showed synaptonemal complex (SC), a meiosis-specific chromosome structure as well as nuclear microtubules from Spindle pole body (SPB), a yeast centrosome embedded in NE ^30–32^. However, it still remains less known about ultra-structures of cellular components inside of meiotic yeast cells.

In this study, we analyzed ultra-structures inside meiotic yeast cells, by using freeze-substitution electron microscope, which is suitable to observe near-native cellular structures ^33^. Interestingly, we detected bundles/cables inside nuclei in cells undergoing meiosis I, but not in pre-meiotic cells or spores. The nuclear bundle is structurally similar to the cytoplasmic bundles induced in meiotic cells. The nuclear and cytoplasmic bundles are immuno-labeled with antiactin antibody and are sensitive to the treatment with an actin-depolymerizing drug. Meiosis-specific nuclear bundles consist of multiple filaments with a regular arrangement forming three-dimensional structure. These results indicate that bundles containing actin or bundle-based networks are formed inside nuclei of cells undergoing unique chromosomal events in prophase I. Moreover, the nuclear bundle is seen in the *spo11* mutant defective in the formation of meiotic DSBs for homologous recombination. The biological implications of nuclear bundles containing actin in meiotic cells are discussed.

## Results

### Electron microscopic observation of meiotic yeast cells

To get more detailed ultra-structures of cellular structures and its spatial relationship with organelles inside of meiotic yeast cells, we used a transmission electron microscope (TEM). Meiosis was induced by incubating yeast diploid cells in sporulation medium (SPM). Under this condition, wild-type cells carried out DNA replication and meiotic recombination from 2 h to 5 h after the induction. At ~5 h, cells entered meiosis I and, by ~ 8 h, most of cells finished meiosis II with further developmental stage of sporulation (Fig. 1). Cells were quickly frozen, and substituted with the fixative and stained with osmium (freeze-substitution method) ^33^. Thin sections of cells (50~60nm) were observed under TEM (Fig. 2). With the freeze-substitution method, cellular organelles including nucleus, mitochondrion, and vacuoles in cytoplasm filled with dense-stained ribosomes were well preserved (Fig. 2a,c,f,j). The nucleus was surrounded with double-layered nuclear membranes and contained electron-dense regions, which corresponds with nucleolus (Fig. 2a,b,c). During meiosis prophase-I, i.e. 4 h after the induction of meiosis, nuclei contact vacuoles forming nuclear-vacuole junctions (NVJ; Fig. 2f) as shown previously ^34^. At 8 h, four pre-spore cells were formed inside of the cells (Fig. 2j).

**Figure 1.**
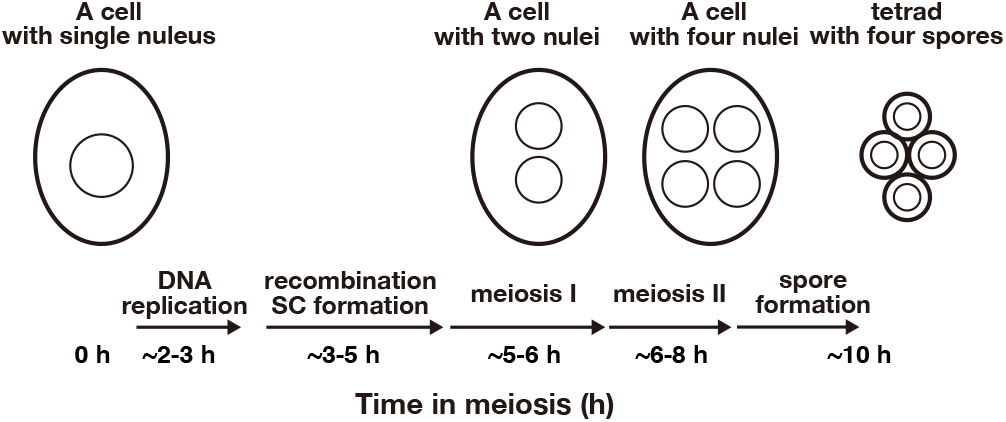
Schematic figure of yeast meiosis.

**Figure 2.**
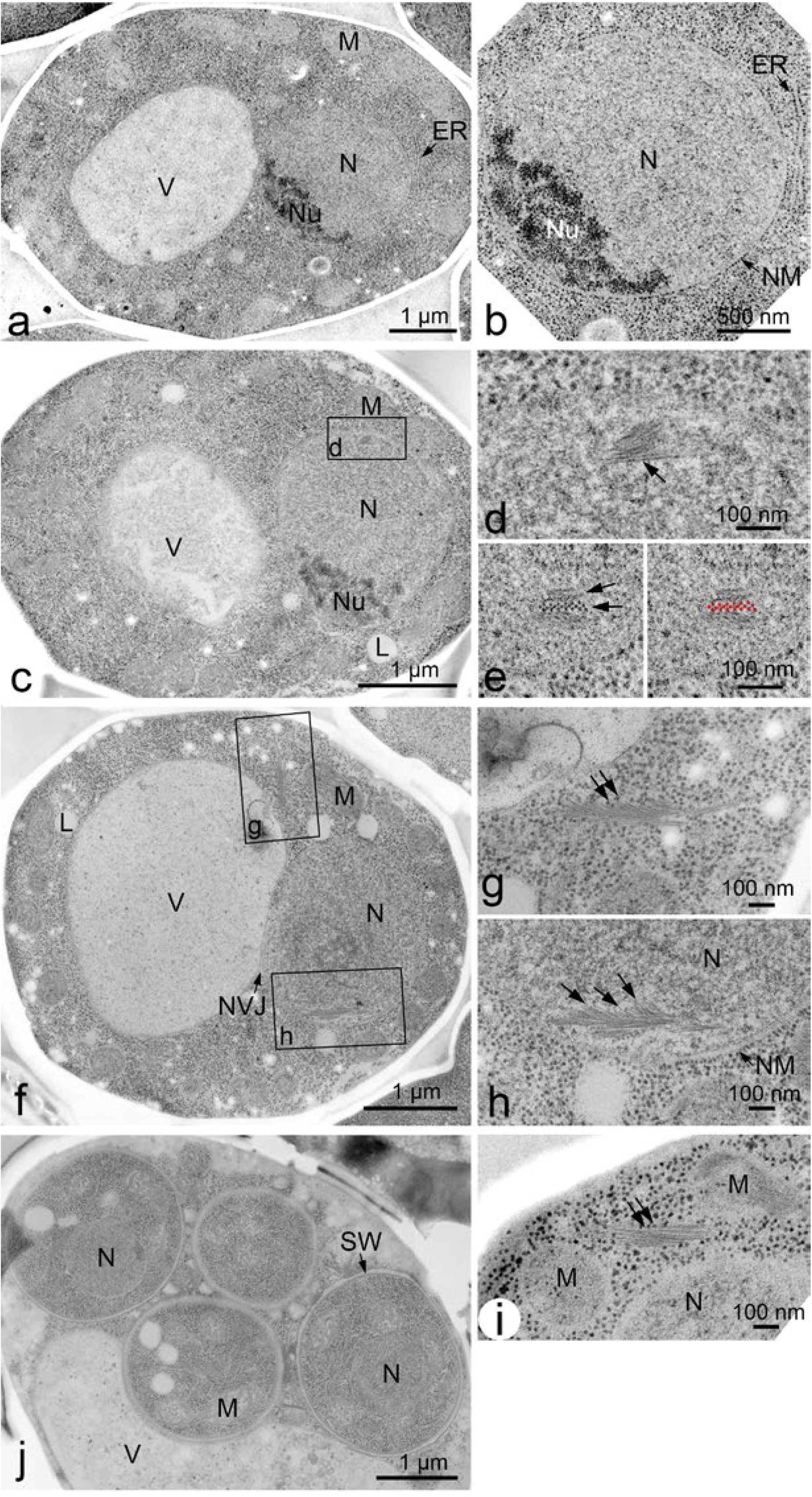
EM images of meiotic yeast cells. **a, b** TEM images of a yeast diploid cell at 0 h. The specimens were prepared with freeze-fixation and sectioned. Magnified image is shown in (b). Bars indicate 1 μm and 500 nm in (a) and (b), respectively. **c-e.** TEM images of a yeast diploid cell at 2 h after incubation with SPM. Magnified views are shown in (d, e). (d) vertical-sectioned of bundles are shown by an arrow. (e) cross-sections of bundles are shown by arrows (left) and marked by red dots (right). Bars indicate 1 μm (c) and 100 nm (d, e). **f-i.** TEM images of a yeast diploid cell at 4 h after incubation with SPM. A whole cell (f) and magnified views of boxed region with bundles in cytoplasm (g) and in nucleus (h), where bundles of feather-like filaments (arrows) are seen. A bundle (arrow) is seen near mitochondrion (i). Bars indicate 1 μm (f) and 100 nm (g, h and i). **j.** TEM image of a yeast diploid cell at 8 h after incubation with SPM. Bars indicate 1 μm. ER, endoplasmic reticulum; L, Lipid body; M, mitochondrion; N, nucleus; NM, nuclear membrane (envelope); Nu, nucleolus; NVJ, nuclear-vacuole junction; SW, spore wall; V, vacuole.

### Nuclear bundles/cables in yeast meiosis

In addition to known cellular structures/organelles, we detected a unique structure of cables or bundles inside of a nucleus of meiotic cells (Fig. 2c, d, e, f, h). This bundle is structurally different from microtubules in the nucleus emanating from SPB (Fig. 3a, b). Sections of meiotic nuclei at late time points such as 5 h contained several bundles (Fig. 3a and Fig. 5), indicating that nuclear bundles are an abundant structure. The bundles contain three to ten thin parallel filaments (Fig. 2e, h). Cross-section of the bundle showed oval-like appearance of a single filament (Fig. 2e and Supplementary Fig. S1). The diameter of a single thin filament in nuclear bundles is around 7-8 nm (Fig. 4a; 8.3±1.3 nm [mean±S.D.; n=17], 7.1±1.1 nm [n=8], 7.6±1.0 nm [n=13] for three independent sections; Supplementary Fig. 1).

**Figure 3.**
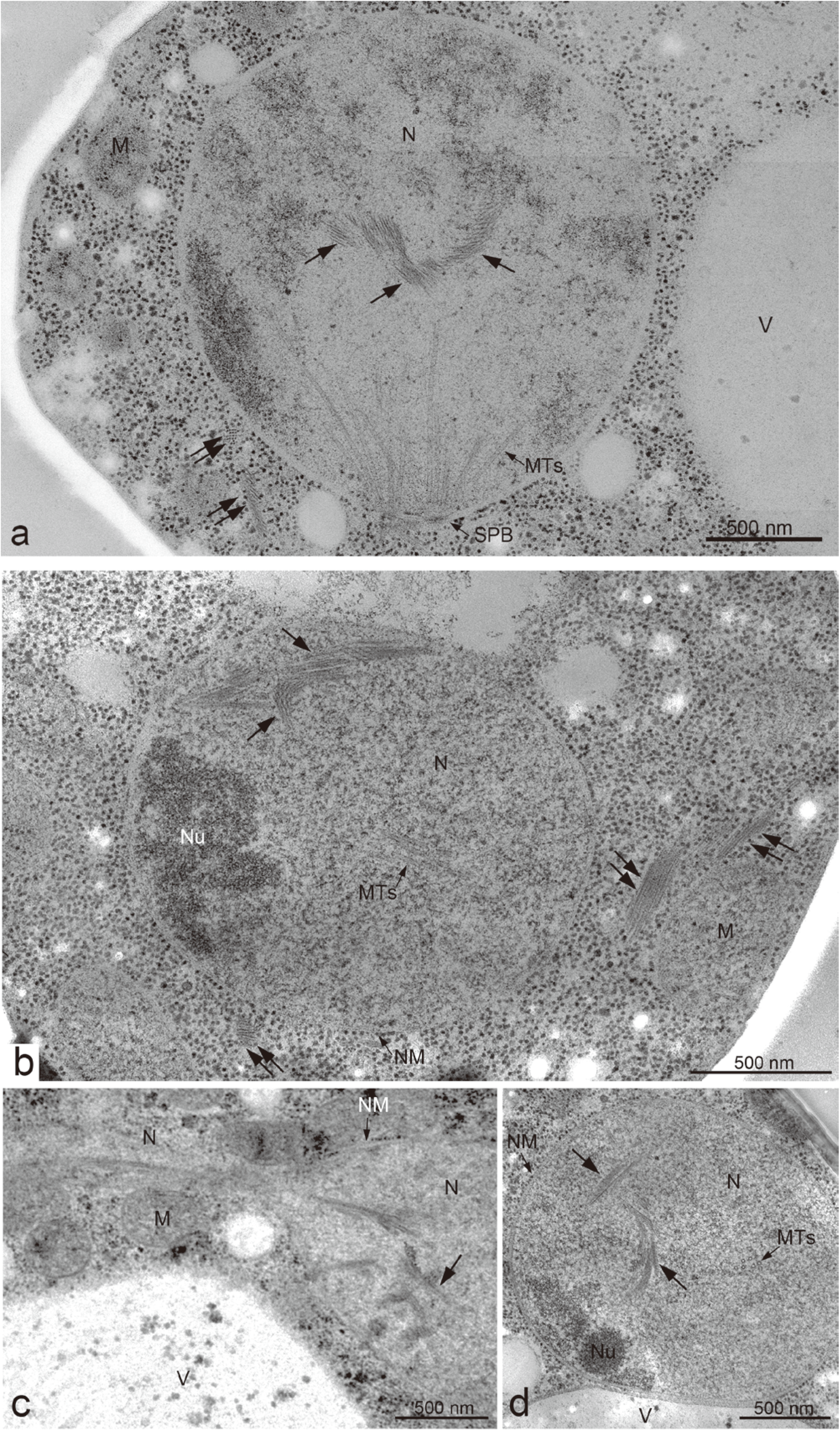
EM images of meiotic yeast cells during mid-prophase I. **a-d.** TEM images of a yeast diploid cell after incubation with SPM for 4 h (a), 5h (b) and 6 h (c, d). The specimens were prepared by cryofixation, freeze-substitution, and sectioned. **a, b, and d,** Representative images of nuclear actin cables with possible branches. **c,** Microtubules and nuclear bundles are found in the stretched nucleus. Black arrows indicate nucleus bundles. Double black arrows indicate cytoplasmic bundles. Bars indicate 500 nm. M, mitochondrion; MTs, microtubules; N, nucleus; NM, nuclear membrane (envelope); Nu, nucleolus; SPB, spindle pole body; V, vacuole.

**Figure 4.**
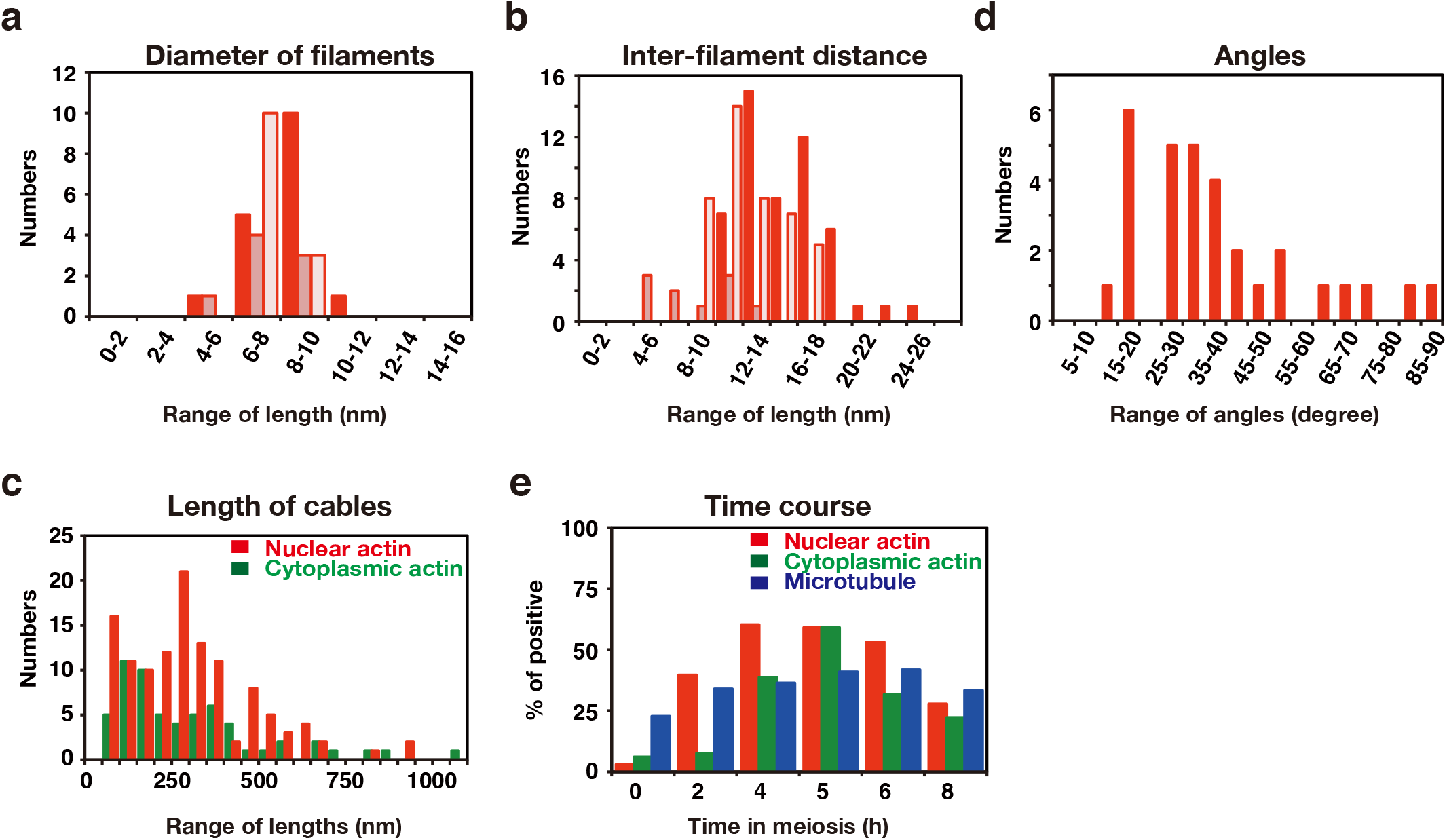
Quantification of nuclear and cytoplasmic bundles during meiosis. **a.** Distribution of a diameter of filaments in nuclear bundles at 4 h in meiosis. The diameter of filaments in a cross-section of nuclear bundles were measured and ranked every 2 nm. The number of each rank in three independent time courses is shown in three different colors. **b.** Distribution of a distance between two adjacent filaments in nuclei at 4 h in meiosis. The distance between two adjacent actin filaments in a cross-section were measured as shown in Supplementary Fig. 1, and ranked every 2 nm. The number of each rank in three independent images is shown in different colors. **c.** Distribution of angles of actin branches in nuclei at 4 h in meiosis. An angle between cables and branched cables was measured as shown in Supplementary Fig. 1. The angles were ranked every 5 degrees. The number of each rank is shown. **d.** Distribution of lengths of actin bundles in nuclei and cytoplasm at 4 h in meiosis. The length of actin bundles in a cross-section were measured and ranked every 50 nm. The number of bundles in nuclei (red) and cytoplasm (green) in each rank in three independent time courses is shown. **e.** Kinetics of the formation of actin bundles in nuclei (red) and cytoplasm (green) as well as nuclear microtubules (blue) at different times of meiosis. A number of sections containing bundles were counted and percentage of actin-positive nuclei and −cytoplasm section as well as microtubule-positive nucleus section are shown.

**Figure 5.**
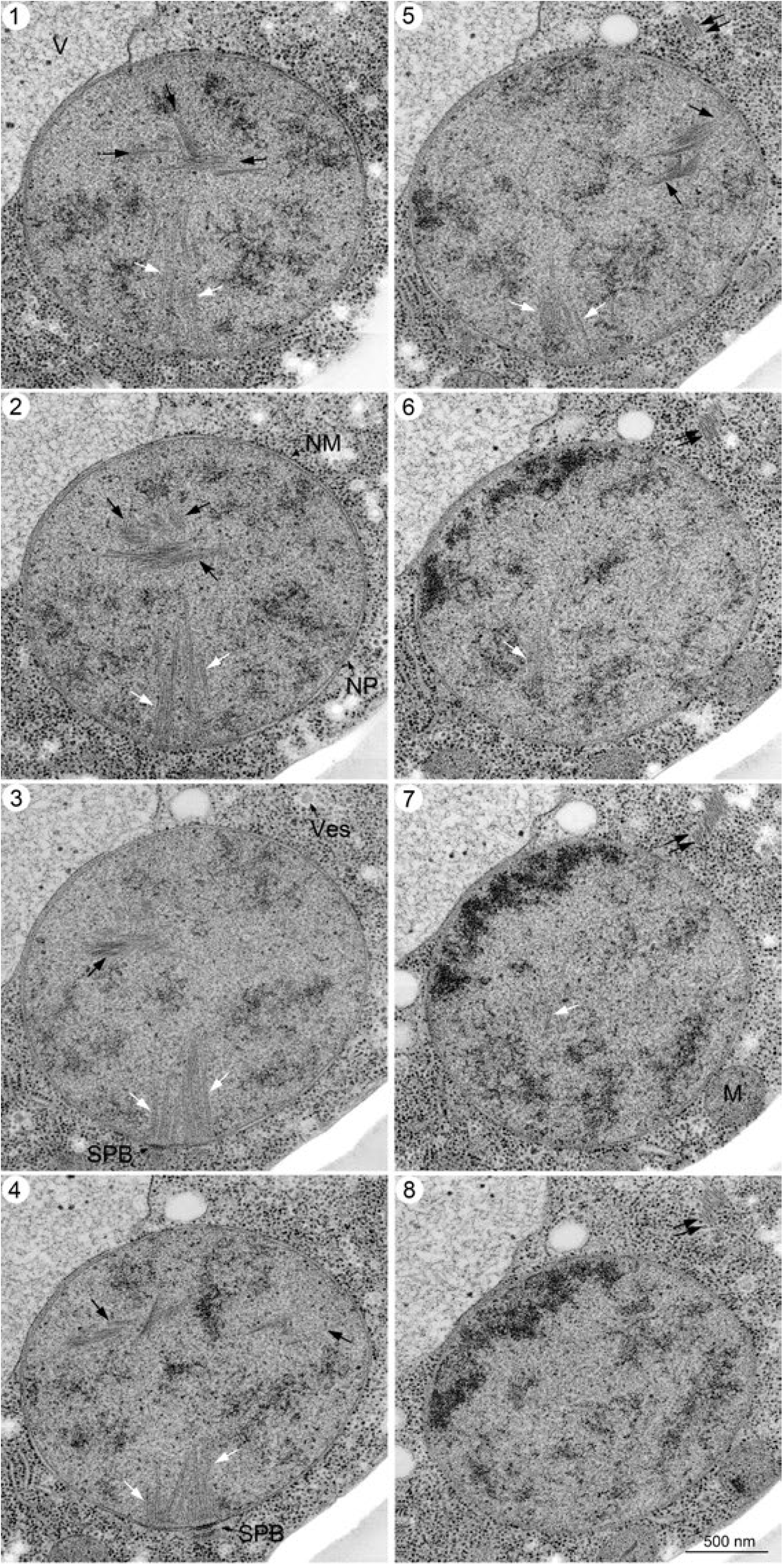
EM images of serial sections of yeast nucleus during midprophase I. Specimens were prepared by high-pressure freezing as described in Materials and Methods. Serial sectioned TEM images (60nm section thickness) of a single cell at 4 h are shown. White and black arrows indicate spindle microtubules and nuclear bundles, respectively. Double black arrows indicate cytoplasmic bundles. Bar indicates 500 nm. M, mitochondrion; NM, nuclear membrane; NP, nuclear pore; SPB, spindle pole body; V, vacuole; Ves, Vesicle.

In the cables, the filaments exhibited a regular rectangular/square arrangement (lattice) with repeated units of alternate single and double filaments (Fig. 2e; shown as red dots in Fig. 2e’). An average distance between adjacent filaments (Supplementary Fig. 1) is 8-15nm (Fig. 4b; 15.2±3.1 nm [n=50], 8.3±3.2 nm [n=10], 12.2±2.5 nm [n=42]). The length of the bundles varies from 50 nm up to 1,000 nm, near the diameter of the nucleus (Fig. 4c). The length of nuclear bundles in sections is 306±223 nm [n=60], median=274 nm (Fig. 4c). In some cases, nuclear bundles span across through whole nucleus (~1 μm, see also Fig. 6-section 5). The longer bundles consist of multiple short bundles, rather than a single linear bundle (see Fig. 6-section 5).

**Figure 6.**
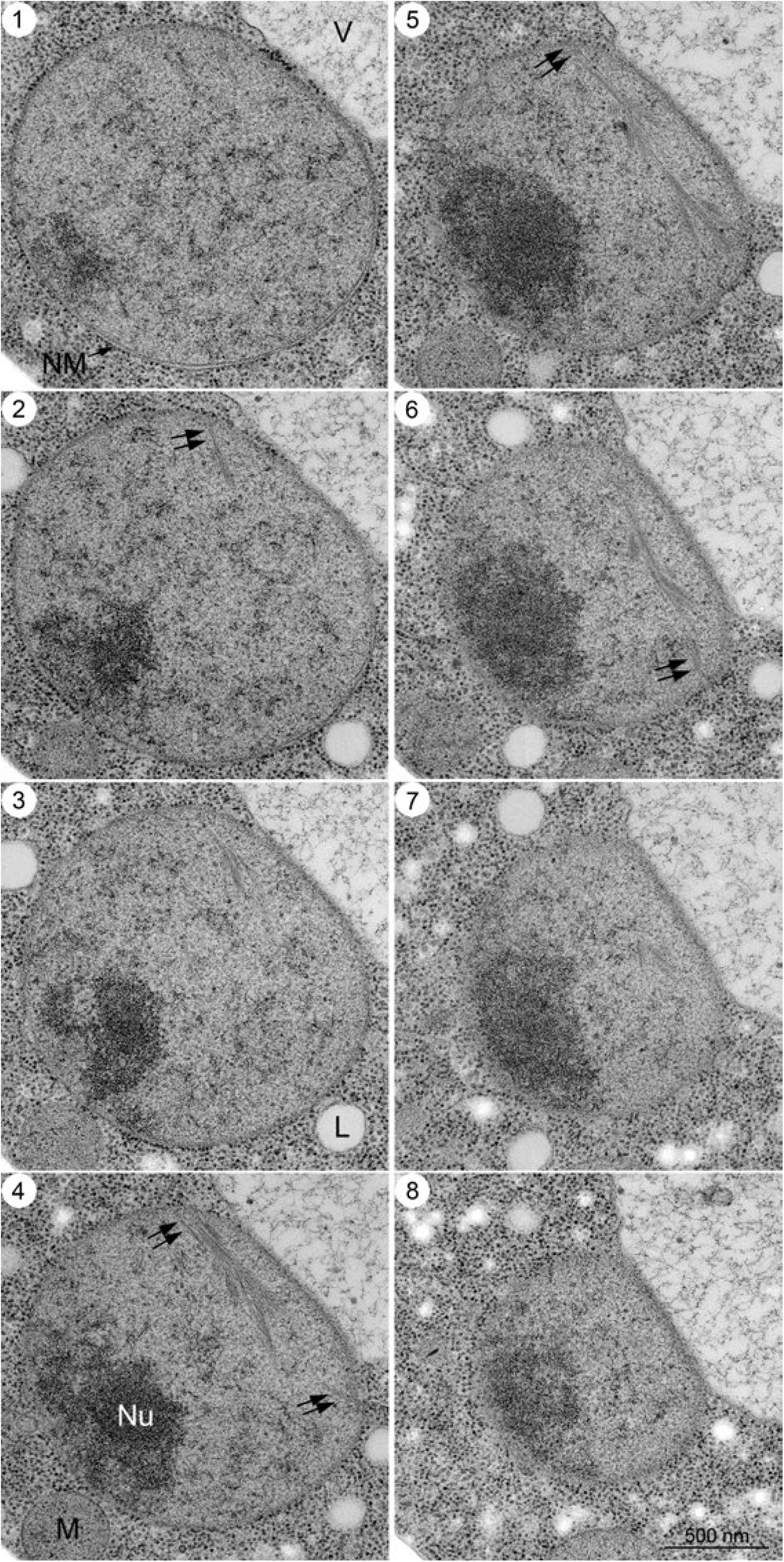
EM images of serial sections of yeast nucleus during mid-prophase I. Serial sectioned TEM images (60nm section thickness) of a cell at 4 h, which were fixed with high-pressure freezing, are shown. Black double arrows indicate bundles, which is likely to attach to nuclear envelop. Bar indicates 500 nm. M, mitochondrion; NM, nuclear membrane; Nu, nucleolus; V, vacuole.

We note a recent preprint by Gan’s group which, by using cryo-electron tomography, showed a nuclear bundle in meiotic yeast cells, which referred to as “meiotic triple helix” (Ma et al. BioRxiv, 10.1101/746982v1). The nuclear bundle found here is structurally similar to the meiotic triple helix.

### Spatial arrangement of nuclear bundles/cables in meiosis

To get more spatial information on nuclear bundles, we checked serial sections of a nucleus in yeast cells at 4 h in meiosis (Fig. 5 and 6). To achieve in-depth freezing in specimens, we froze cells under a high pressure ^35^. With high-pressure freezing specimens suitable for sectioning were obtained. Importantly, we did not see any change of sub-cellular structures including cables in yeast cells prepared by either rapid freezing or high-pressure freezing (compare Fig. 2 with Fig. 5 and 6). We often detected multiple nuclear bundles in different sections of 60 nm (Fig. 5, 6), indicating that the cables are an abundant nuclear structure with three-dimensional arrangement of cables in a nucleus (Fig. 5). In some sections, a long bundle that spans entire nucleus is observed (Fig. 6-sections 5 and 6), and the end of the cable is likely to attach to the NE (arrows in Fig. 6-section 4-6).

Nuclear bundles often accommodate branched-like filaments or bundles, which look like “feather” (Fig. 2g, h). At this resolution some branched filaments look attached a lateral side of filaments in a main bundle while the other filaments are not attached. We measured an angle between main bundles and branched bundles/filaments (Fig. 4d, Supplementary Fig. 1). At 4 h, angles between the main bundle and branch-like filaments are 25°-40° with sub-peak of 15°-20° (Fig. 4d; 38°±19° [n=31], median=34°). We noticed that branch-like filaments from a single bundle are oriented same direction, suggesting the presence of the directionality of the cable.

### Nuclear bundles are similar to cytoplasmic bundles in meiosis

We also detected cables/bundles in cytoplasm of meiotic yeast cells (Fig. 2g, i). Structurally, the bundles in cytoplasm are similar to those in nuclei (compare Fig. 2g, h). The diameter of a single thin filament in cytoplasmic bundles is around 7-8 nm (Supplementary Fig. 2a; 7.1±1.0 nm [n=21]), which is similar to that of nuclear bundles (Fig. 4a). The length of cytoplasmic bundles in sections (mean=304±179 nm [n=121], median=238 nm; Fig. 4c) is not different from that in cytoplasm (*P*=0.64, Mann-Whitney’s *U*-test). These indicate that nuclear and cytoplasmic bundles/cables are structurally indistinguishable. While nuclear bundles form complicated 3D structure, cytoplasmic bundles form less such the structure (Fig. 3b).

### Nuclear bundles by chemical fixation

Yeast cells are surrounded with thick cell walls, which impair penetration of staining reagents such as osmic tetroxide. We also stained spheroplasts of meiotic yeast cells with osmic acid after fixation with glutaraldehyde (without any freezing, Supplementary Fig. 3) and, in some cases, prepared serial sections for EM observation (Supplementary Fig. 4). With this procedure, the mitochondria are contrasted highly (Supplementary Fig. 3). Membrane structure such as nuclear membrane was partially deformed, possibly due to hypo-osmotic conditions. Importantly, even under this condition, we could detect the cables/bundles in both the nucleus and the cytoplasm of cells during meiotic prophase-I, whose structures and arrangement are similar to those obtained by the freeze-substitution method.

### Nuclear and cytoplasmic bundles are formed during prophase-I

We checked the presence of nuclear bundles in different stages of meiosis (Fig. 4e). As a control, we measured nuclei containing microtubules. The nuclear microtubules are seen at 0 h and sections positive to microtubules are increased slightly during meiotic prophase-I. Nuclear bundles appear earlier during meiosis than cytoplasmic bundles (Fig. 4e). At 0 h before the induction of meiosis in which most of cells are G1, we detected few bundles in nuclei (Fig. 2a; 1.9% [0/10, 1/10 and 1/34, numbers of bundle-positive nucleus section/numbers of total nucleus section examined, three independent time courses) as well as in cytoplasm (4.5% [1/10, 1/22 and 1/34] numbers of bundle-positive cytoplasm section/numbers of total cytoplasm section examined, three independent time courses). Few bundles were also found in four nuclei of pre-spore cells (Fig. 2j), suggesting that the bundles disassemble by the formation of pre-spore cells. Kinetic analysis revealed that nuclear bundles were seen at 2 h post-induction of meiosis (Fig. 4e), when 38.9% (13/31, 8/22) of sectioned cells contained nuclear bundles, indicating the formation of the structure is an early event of meiosis; e.g. during pre-meiotic S-phase. At 4 h, cable-positive nuclei were increased to ~60% frequency (1/12, 12/22, 15/22, 25/32; numbers of bundle-positive nucleus section/numbers of total nucleus section examined, four independent time courses). Cells in late prophase-I at 5 h showed a peak in both bundle-positive nuclei and cytoplasm (Fig. 4e). After 6 h both nuclear and cytoplasmic bundles were decreased (Fig. 4e). At 6 h, when about a half of the cells enter in meiosis I, we detected the bundles in a bridge-like structure between two nuclei undergoing anaphase-I (Fig. 3c). This suggests that some nuclear bundles passed into the daughter nuclei during meiosis I division.

### Nuclear bundles are formed in the absence of meiotic DSBs

Since nuclear bundles are induced during meiotic prophase I, we analyzed the relationship of the bundle with other prophase I events such as meiotic recombination and SC assembly, both of which depends on the formation of meiotic DSBs by Spo11. We examined the bundle formation in the *spo11-Y135F* mutant, which is defective in meiotic DSB formation ^36, 37^. At 0 h, there is little bundles in both cytoplasm and nucleus (Fig. 7a). As in wild-type cells, at 4 h, the bundles are found in both cytoplasm and nucleus in the *spo11-Y135F* mutant cells (41.2%, Fig. 7b). The kinetics of nuclear and cytoplasmic bundles (Fig. 7c, d) in the *spo11* mutant are similar to those in wild type (Supplementary Fig. 5). These indicate that the formation of nuclear bundles as well as cytoplasmic bundles are independent of meiotic DSBs induced by Spo11, thus meiotic recombination and the SC.

**Figure 7.**
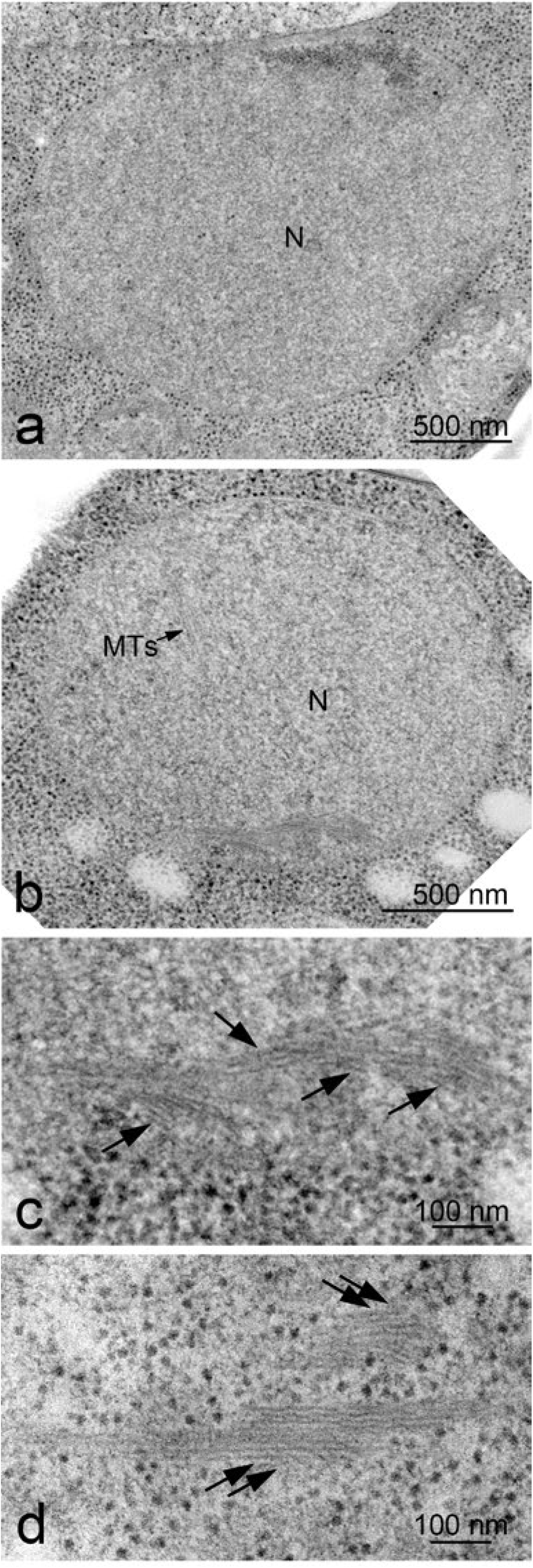
Nuclear bundles formed in the *spo11* mutant. TEM images of a yeast *spo11-Y135F* diploid cell at 0 h (a) and 4 h (b). The specimens were prepared with freeze-fixation and sectioned. Magnified images are shown in (c, d). Branch-like structure is shown in arrows. M, mitochondrion; N, nucleus; MTs, microtubules. Bars indicate 500 nm (a, b) and 100 nm (c, d).

### Nuclear and cytoplasmic bundles contain actin

Previous EM analyses of mitotic cells of both budding and fission yeasts have shown three distinct actin sub-cellular structures; rings, cables, and patches as well as less-defined filasome ^21–24^. As a less-defined actin-related structure, filasome is a cytoplasmic structure of less electron-dense areas with a vesicle in the center (Supplementary Fig. 6), a novel actin-containing membrane-less sub-cellular structure in cytoplasm originally found in fission yeast ^24^.

We initially found that EM images of meiotic nuclear and cytoplasmic bundles are similar to those of actin cables in yeasts ^21–25^. In addition, live visualization of cytoplasmic actin bundles in meiotic prophase I with Abp140-GFP, which marks actin cables ^27, 28^, indicated the presence of multiple actin cables only in cytoplasm.

It is known that the treatment of meiotic cells with an actin-depolymerizing chemical, Latrunculin B (LatB), largely reduced cytoplasmic actin cables ^27, 28^. To confirm that the bundles in nuclei as well as in cytoplasm are indeed dependent of actin polymerization, we treated 4-h meiotic yeast cells with an actin-depolymerizing drug, Latrunculin B (LatB), for 1 h and examined actin cables in cells under EM. As shown in Fig. 8a, b, the number of bundles (a section positive to actin bundles) is mildly reduced in nuclei after the treatment with LatB (from 60% [1/12, 12/22, 15/22, 25/32] at 4 h for positive nucleus to 31% [8/14, 3/21]) after 1 h-treatment; *P*=0.0247, Fisher’s exact test) without affecting nuclear microtubules. This indicates that some nuclear bundles are sensitive to LatB, supporting the idea that bundles both in nucleus and cytoplasm are formed through actin polymerization. Consistent with this, the bundle is also sensitive to the other actin-depolymerizing chemical, Latrunculin A (LatA) (Ma et al, preprint).

**Figure 8.**
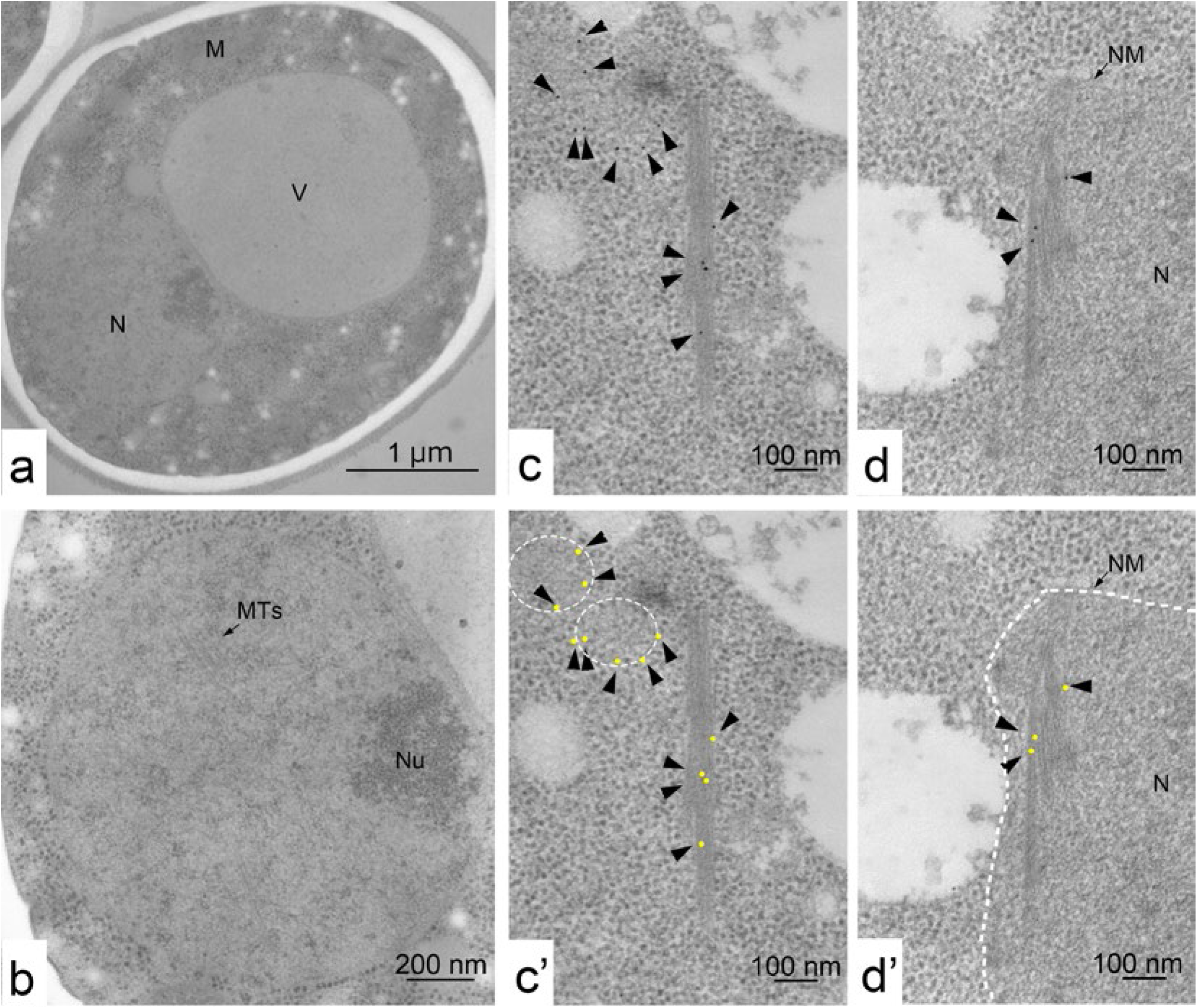
Nuclear bundles contain actin and its formation depends on actin polymerization. **a, b.** TEM image of a Latrunculin B-treated cell (a) and nucleus (b). Cells were incubated with SPM for 4 h and treated with Latrunculin B for 1 h. The specimens were prepared by rapid-freeze with fixation and sectioned. Bars indicate 1 μm (c) and 200 nm 8b). M, mitochondrion; N, nucleus; NM, nuclear membrane; Nu, nucleolus; MTs, microtubules; V, vacuole. **c, d. I**mmuno-gold labeling using anti-actin antibody was carried out as described in Materials and Methods. Images of cables in cytoplasm (c) and in nucleus (d) in a cell at 4 h are shown. The positions of gold particles are shown in arrowheads. Dotted circles were actin patches or filasomes (c’). The area surrounded by dotted line shows a nucleus (d’). The positions of gold particles are marked by yellow dots (c’ and d’). Bars indicate 100 nm.

Previously, immuno-EM confirmed the presence of actin in the bundles in cytoplasm of yeasts ^38^. We performed immuno-gold labeling of chemical fixation sections using anti-actin antibody, which provides less clear image compared to conventional EM (Fig. 8c, d). Although not extensive, we found clustered gold labels on the bundles in nuclei and in cytoplasm (Fig. 8c, d). In a nucleus, in addition to the bundles, we often detected the particles on dense-stained area containing filament-like structure, which might be bundles. Importantly, nuclear bundles of freeze fixation specimens showed more gold particles than other nuclear area (Supplementary Fig. 7). These suggest that both nuclear and cytoplasmic bundles contain actin. We also detected gold particles on less electron-dense areas in cytoplasm without ribosomes, which might correspond to the filasome (Fig. 8c).

## Discussion

In this study, by using TEM with rapid freezing-fixation, we found nuclear bundles/cables containing actin in budding yeast cells undergoing the physiological program of meiosis. We could also detect the bundles in cytoplasm, which are also induced during prophase I. Since we used rapid freezing to preserve structures inside of cells, it is unlikely that the cable is an artifact produced by specimen preparation, which might be induced by external stress and/or staining. Moreover, we also detected the bundles in nuclei fixed with chemicals without freezing (Supplementary Fig. 3 and 4). We also noted that cryo-electron tomography showed a nuclear bundle in meiotic yeast nuclei, which is similar to the nuclear bundle described in this paper (Ma et al. BioRxiv, 10.1101/746982v1).

Both nuclear and cytoplasmic bundles are structurally similar, consisting of multiple parallel filaments (Fig. 9a). Nuclear (and cytoplasmic) bundles seem to contain a unique arrangement of filaments, in which alternate pattern of 1 and 2 filaments is observed in a cross section of the cables (Fig. 9b). This alternate pattern provides rectangular/square arrangement of filaments in a single bundle. The distance of inter-filaments is mainly 10-15 nm. Although we sometimes observed branch-like structure of the bundle, we have not had any solid evidence on the branching of the filaments at current resolution of EM.

**Figure 9.**
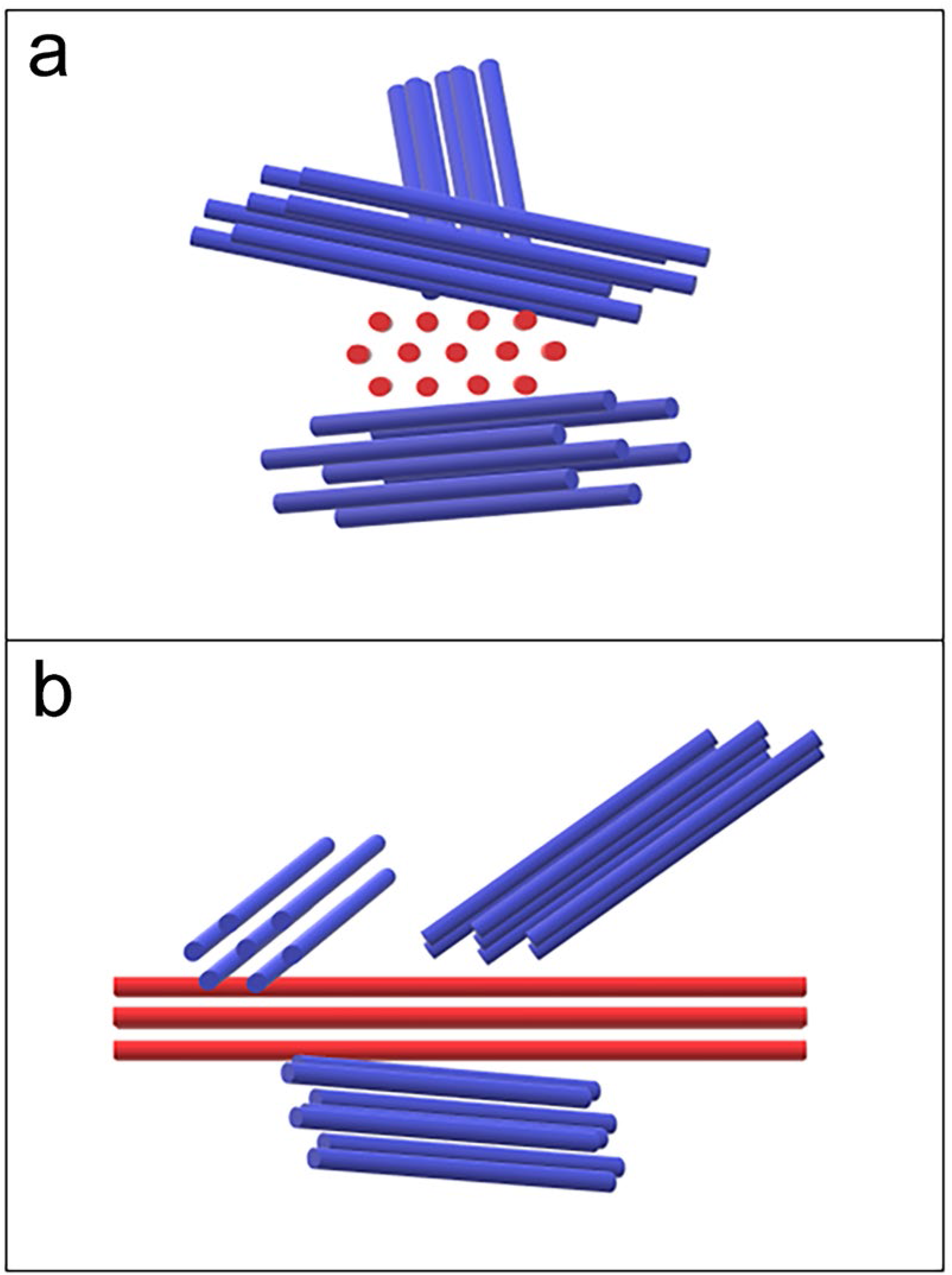
A model of nuclear bundles. A schematic representation of actin bundles (red) with multiple branch-like filament (blue) is shown. Schematic presentation of a cross-section of a nuclear actin bundle (a); arrangement of actin filaments in the bundle (b).

Nuclear bundles elongate up to 1 μm (Fig. 4c and Fig. 6). This long bundle consists of several bundles in a linear array, rather than a single bundle, suggesting a self-assembly property of the bundle. We speculate that nuclear bundles are assembled into a long cable with lateral attachment (or branching). Several bundles are present in a single nucleus of meiotic cells (Fig. 5 and 6), indicating that nuclear bundles are abundant with three-dimensional arrangement during late prophase I. The nuclear bundles formed during meiosis appear to have a unique ultra-structure: multiple bundles/cables accommodate threedimensional arrangement, possibly through lateral interaction among the bundles and/or branch-like configuration. If abundant, how these cables are packed in the context of nucleoplasm with meiotic chromosome structures such as SCs remains to be determined. Unfortunately, we could not efficiently detect SCs in our cryo-sections, which could be detected in silver-staining of fixed meiotic cells^30–32^.

Nuclear bundles are induced from very early meiotic prophase-I such as 2-h post induction of meiosis and are present to at least by meiosis I nuclear division. The bundles are abundant not only in nuclei, but also in cytoplasm particularly during late prophase-I (Fig. 3b). On the other hand, we rarely see nuclear or cytoplasmic bundles in a G1 diploid cell (pre-meiotic) or in spores (Fig. 2a, i). During meiosis, the formation of nuclear bundles starts at 2 h post-induction of meiosis, prior to DSB formation, which begins at ~3 h, indicating that nuclear bundle formation is not be associated with DSB formation during yeast meiosis. Indeed, the bundles are formed in the absence of meiotic DSBs in the *spo11* mutant.

Our results here suggest that meiotic nuclear bundles contain actin. Are a filament in nuclear and cytoplasmic bundles described here so-called a typical actin filament with double-helix? The diameter of the filament in nuclear and cytoplasmic bundles is around 7-8 nm at minimum, which roughly corresponds with the size of a single yeast actin filament reconstituted *in vitro* ^39, 40^. However, the angle of putative branching in nuclear bundles is quite different from an angle of branches of actin filament mediated by Arp2/3 complex with a value of 70° ^41^. Previous staining analyses with phalloidin or imaging of actin-probe, Abp140-GFP ^42^ or Life-Act (Ma et al. BioRxiv), have not shown the presence of any actin polymers inside of yeast meiotic nuclei. These suggest that the nuclear bundle/cable is a novel structure containing actin. Indeed, cryo-electron tomography showed the nuclear bundle contains triple-helical filaments (Ma et al. BioRxiv).

What is a role of nuclear bundles? Actin polymerization in cytoplasm is involved in chromosome motion during prophase-I in budding yeast ^42^. Cytoplasmic actin cables promote meiotic chromosome motion through SUN-KASH protein ensembles in the NE ^29^, which is sensitive to the LatB ^27^. Our results here suggest that nuclear bundles also play a role in the movement of meiotic chromosomes. Alternatively, nuclear bundles may protect nuclear structures from external forces generated during meiosis as seen in *Xenopus* oocyte ^16^, by providing a rigid structure that resists mechanical stress generated through chromosome motions. We speculate that nuclear bundles may form threedimensional structure through self-assembly inside of meiotic nuclei. Further studies are necessary to reveal nature and function of nuclear bundles containing actin in meiotic cells.

## Acknowledgements

We thank Drs. N. Nagata and I. Mabuchi for discussion and suggestions. We also thank members of the Laboratory of Electron Microscopy, Japan Women’s University and Ms. Y. Osaki in NPO: Integrated Imaging Research Support. We thank Drs. Gasser, Hurst, Shimada, Mabuchi, Yasunaga, and Usukura for critical reading of the manuscript. This work was supported by JSPS KAKENHI Grant Number: 22125001, 22125002, 15H05973 and 16H04742 to A.S and by the Open Research Center of JWU established in private universities in Japan with the support of the Ministry of Education, Culture, Sports, Science and Technology to M. O.

## Author contributions

TT, MO, and AS conceived and designed the experiments. TT performed the all EM experiments. TT and AS analyzed the data. TT, MO, and AS prepared the manuscript.

## Materials and Methods

### Strains and culture

*S. cerevisiae* SK1 diploid strain, MSY832/833 (*MATα/MAT**a**, ho::LYS2/”, lys2/”, ura3/”, leu2::hisG/”, trp1::hisG/”*) and HSY185/186 (*MATα/MAT**a**, ho::LYS2/”, lys2/”, ura3/”, leu2::hisG/”, spo11-Y135F-KanMX6/”*) was used for meiotic time course.

Yeast cell culture and time-course analyses of the events during meiosis and the cell cycle progression were performed as described previously ^43, 44^. Briefly, 1 ml of diploid yeast culture in YPAD (1% yeast extract, 2% Bacto peptone, 2% glucose) was cultured in 200 ml of SPS (0.5% yeast extract, 1% Bacto peptone, 0.17% yeast nitrogen base, 1% KCH_3_COO, 1% potassium hydrogen phthalate) media for 16 h. Cells were collected and, after washing with H2O, were re-suspended in 200 ml of SPM media (0.3% KCH_3_COO, 0.02% raffinose) to start meiosis.

### Rapid Freezing for transmission electron microscopy

Specimens for freeze-substitution electron microscopy were prepared according to previously described method ^45, 46^, with slight modifications. Cells were harvested by centrifugation. The cell pellets were sandwiched between two copper disks (3 mm in diameter). Specimens were quickly frozen with liquid propane using a rapid freezing device (KF80; Leica, Vienna, Austria). Specimens were freeze-substituted in cold absolute acetone containing 2% osmium tetroxide (OsO_4_) at –80°C for 48-72 h and were then warmed gradually (at −40°C for 4 h, at −20°C for 2 h, at 4°C for 2 h and at room temperature for 2 h) and washed with absolute acetone and rinsed with QY-2. Substitution for embedding was infiltrated with Quetol-812 mixture (10, 30, 50, 70, 80 and 90 and 100% pure resin). The specimens polymerized at 60°C for 2 days.

### High pressure freezing fixation for transmission electron microscopy

The specimens for high-pressure freeze-substitution electron microscopy were prepared according to previously described method ^47^ with slight modifications. The pelleted cells were pipetted into aluminum specimen carriers (Leica) and frozen in a high-pressure freezing machine HPM-010 (BAL-TEC, Liechtenstein). The cells were transferred to 2% OsO_4_ in cold absolute acetone. Substitution fixation was carried out at −90°C for over 80 h. After the fixation, the specimens were warmed gradually (at −40°C for 2 h, at −20°C for 2 h, at 4°C for 2 h and at room temperature for 1 h) and washed with absolute acetone and then with exchange to 0.1% uranyl acetate in absolute acetone. After the staining, specimens were washed with absolute acetone and rinsed with QY-2. Substitution and embedding were described in above.

### EM grid preparation and observation

Specimens for morphological observation were sectioned using a ULTRACUT-S ultramicrotome (Reichert-Nissei, Tokyo, Japan). Ultrathin sections were cut with thickness of 50-60 nm and mounted on copper grids. Specimens for immuno-staining were mounted on nickel grids. Ultrathin sections were stained with 4% uranyl acetate for 12 min in dark at room temperature and citrate mixture (SIGMA-ALDRICH) for 2 min at room temperature, and examined using JEM-1200EXS at 120kV or JEM-1400 transmission electron microscope (JEOL, Tokyo, Japan) at 100 kV.

### Immunoelectron Microscopy and Immunolabelling

For immuno-electron microscopy, chemical fixation specimens shown below were etched with 1% H2O2 for 5min at room temperature. Otherwise, cell pellets were sandwiched between two aluminum disks (diameter 3 mm). And specimens were quickly frozen with liquid propane. They were freeze-substituted in cold absolute acetone containing 0.01% OsO_4_. In other ways, specimens were fixed with high pressure freezing machine. They were freeze-substituted in cold absolute acetone containing 0.1% glutaraldehyde. Substitution took place for 48-72 h at −80°C, and were then warmed gradually (at −40°C for 4 h, at −20°C for 2 h and at 4°C for 1h) and washed with dehydration ethanol at 4°C. Then the cells were washed twice with cold ethanol and substituted at 4°C and infiltration was done in the LR white and ethanol mixture (10, 30, 50, 70, 80, 90, 100% pure resin). Specimens were embedded in pure resin and polymerized at 50°C for 1-2 days.

Immunolabelling was carried out by previously described method ^47^ with a slight modification. The sections mounted on nickel grids were incubated with chromatographically purified goat IgG (Zymed Laboratories, Inc., San Francisco, USA) diluted in blocking buffer, 0.1% bovine serum albumin (Sigma Chemical Co., St. Louis, USA) in 50 mM TBS (137 mM NaCl, 2.7 mM KCl, 50 mM Tris-HCl; pH 7.5), to 1/30 concentration for 30 min at room temperature. Affinity-purified mouse anti-actin monoclonal antibodies (MAB1501: Chemicon International Inc., Temecula, USA) diluted in blocking buffer to 1/100 concentration were applied to the sections, incubated for 1 h at room temperature. The sections were washed with blocking buffer. Goat antibodies to mouse IgG labeled with 10 nm colloidal gold (British BioCell International, Cardiff, UK) diluted in blocking buffer to 1/40 concentration and applied to sections for 1 h at room temperature. Sections were washed with blocking buffer and running water. The sections were stained with 4% aqueous uranyl acetate and citrate mixture (Sigma Chemical Co., St. Louis, USA).

### Latrunculin B treatment

After cultured in SPM medium for four hours, cells were treated with 30 μM Latrunculin B (R&D Systems) dissolved in 0.1% DMSO at 30°C for 1 hour. After the post-treatment, cells were collected and fixed by the rapid freezing mentioned later.

### Chemical fixation method for electron microscopy

Cells for chemical fixation were shaking-cultured for 4 hours in SPM containing 1 M sorbitol (SPMS) at 130 rpm and 30°C, details described above. Specimens for chemical fixation were shaking-cultured for 4 hours in SPM containing 1 M sorbitol (SPMS) at 130 rpm 30°C, details described above. Cells were treated for 2 min with 0.2 M DTT in ZK buffer (25 mM Tris-HCl and 0.8 M KCl). Then cells were treated for 30min with 0.5 mg ml^-1^ Zymolyase 100T (Seikagaku Co., Tokyo) in ZK buffer at room temperature. And cells treated with 0.5% Triton X-100 dissolved in PEM (20 mM PIPES, 20 mM MgCl2, 10 mM EGTA, pH 7.0), protease inhibitors, and 1 M sorbitol for permeabilization in the presence of 4.2 μM phalloidin (Sigma-Aldrich) for 1 min, these treatments were performed as described previously ^21, 48^.

Specimens for TEM were prepared as described previously ^24, 48–50^. Briefly, harvested cells were fixed with 2% glutaraldehyde (Electron Microscope Sciences) and 0.2% tannic acid (TAAB) for 1 h at 4°C. After washings with PEM, cells were post-fixed in 2% OsO_4_ in PEM for 1 h at 4°C. After washed out with D.W., cells were embedded in 2% agarose and cut into blocks smaller than 1 mm^3^. And then, specimens were dehydrated with ethanol series (50%, 70%, 80%, 90%, 95%, 99.5%, and Super Dehydrated ethanol). Specimens rinsed with QY-2. Gradually increase the concentration of the resin infiltrated with Quetol-812 (Nisshin EM, Tokyo) mixture (10, 30, 50, 70, 80, 90 and 100% pure resin).

### Image processing and measurement

Measurement of diameter of actin cables and distance between actin cables was carried out using ImageJ Fiji. ^51^. Cropped regions of TEM images were converted into binary images and noises were removed. Filaments more than 10nm^2^ in size and 0.5 - 1.0 in circularity were recognized with analyze particles function. Minor axis lengths of each particle were regarded as diameters of filaments. Distances between filaments were calculated using the central coordinate of particles.

### Statistics

Statistical significance for Length of actin cables between that in nucleus and cytoplasm was analyzed using the Mann-Whitney’s *U*-test. The null hypothesis was that there exists no variation of the length between nucleus and cytoplasm.

Statistical significance for existence of actin cables in cells with and without LatB treatment was analyzed using Fisher’s exact test. The null hypothesis was that LatB treatment did not affect existence of actin cables in cells. Two-sided *P*-value was shown.

## Data availability

Strains are available upon request. The authors affirm that all data necessary for confirming the conclusions of the article are present within the article and figures. EM images are available upon request.

## Supplementary Figures

**Supplementary Figure 1.**
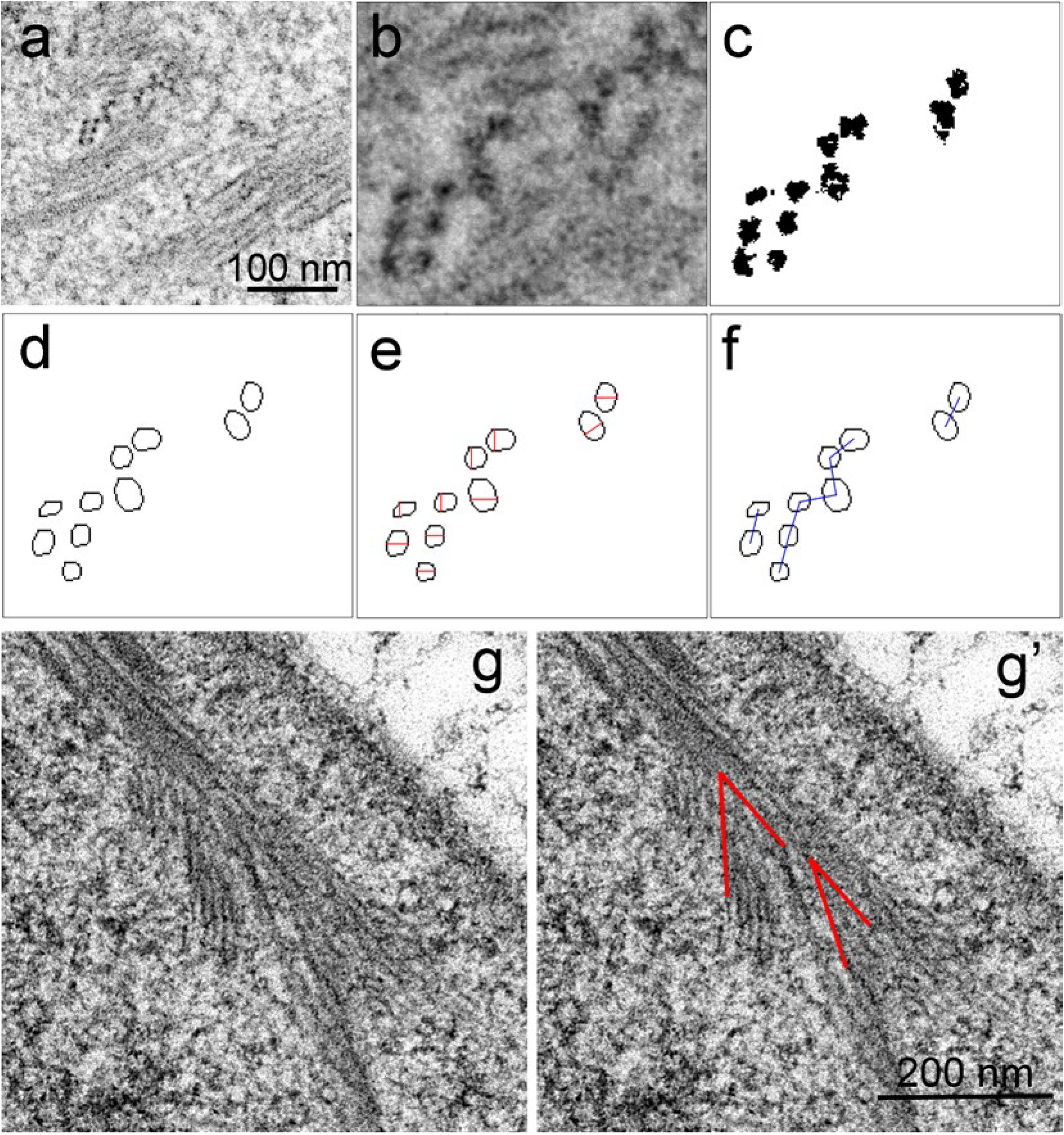
EM images of bundles for measurement of diameter and inter-filament distance. **a-f.** A representative image of cross-sectioned bundles is shown (a) with a magnified view (b), which is converted to a noise removed binary image (c). An elliptical fitting image of each filament was taken (d). A short axis (red) was measured as a diameter (e). Distances between centers of each filament were measured as an inter-filament distance (bule line, f). **g.** Representative images of nuclear bundles with possible branches. Two representative lines (red in g’) were drawn for the measurement for an angle. Bar indicates100nm (a) and 200 nm (g).

**Supplementary Figure 2.**
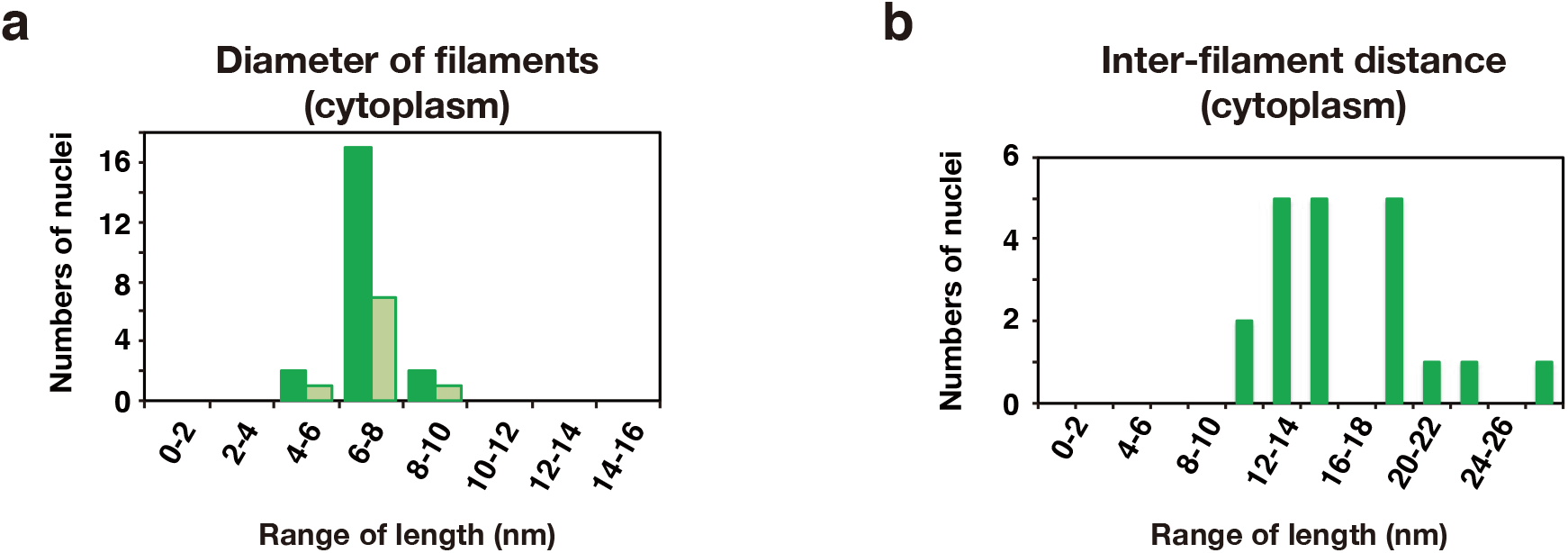
Properties of cytoplasmic bundles. **a.** Distribution of a diameter of the filament in cytoplasmic bundles in nuclei at 4 h in meiosis. The diameter of filaments in a cross-section were measured shown in the main text and ranked every 2 nm. The number of each rank in two independent images is shown in different colors. **b.** Distribution of a distance between two adjacent filaments in cytoplasm at 4 h in meiosis. The distance between two adjacent filaments in a cross-section was measured shown in the main text and ranked every 2 nm.

**Supplementary Figure 3.**
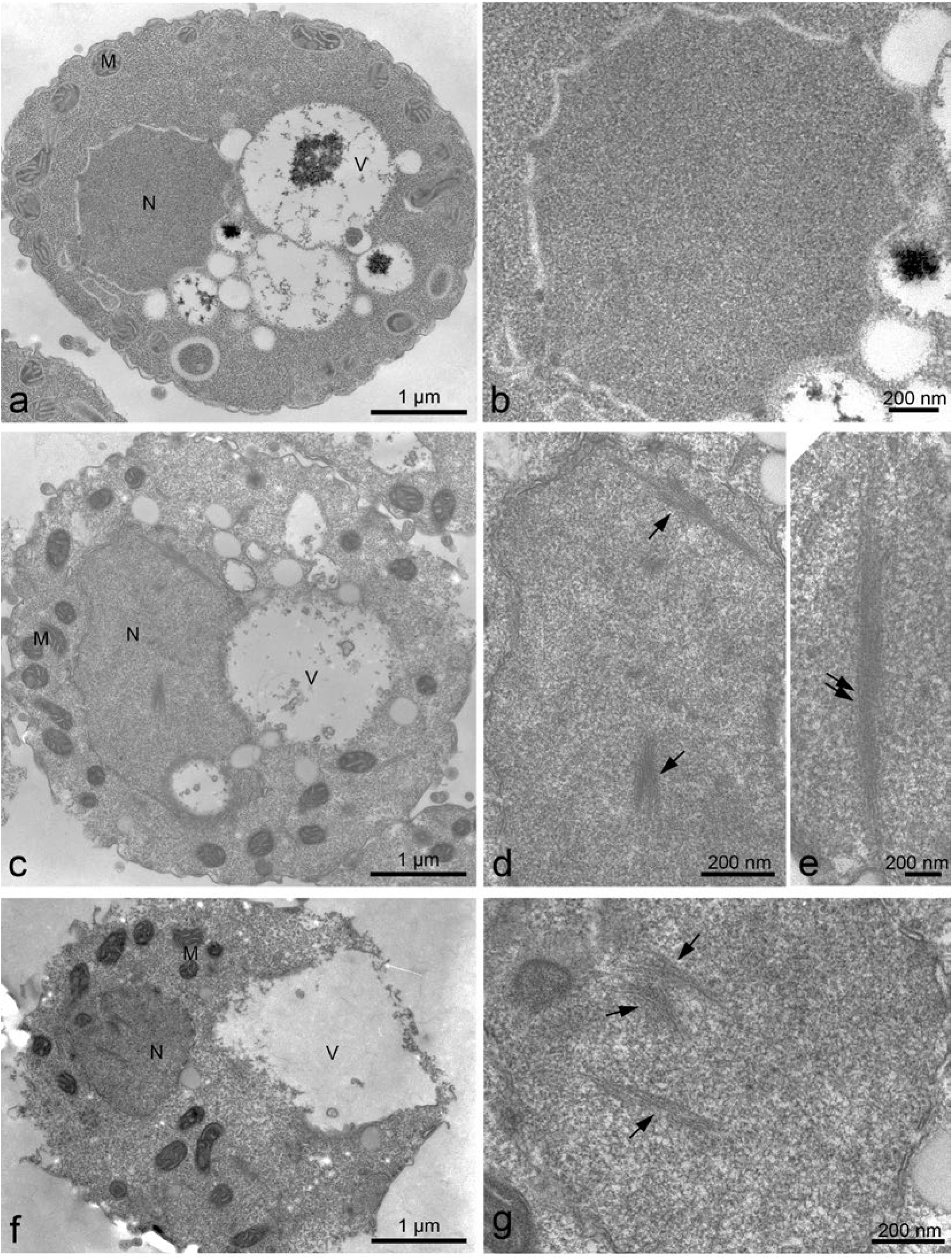
EM images of chemically fixed meiotic yeast cells in mid-prophase I. TEM images of a yeast diploid cell after incubation with SPM for 0 h (a, b), 4 h (c, d, e), and 6 h (f, g). The specimens were fixed with chemicals (glutaraldehyde and OsO_4_) and sectioned. Magnified images with a bundle in nucleus (d, g) and in cytoplasm (e) are shown. Bars indicate 1 μm (a, c f) and 200 nm (b, d, e, g). M, mitochondrion; N, nucleus; V, vacuole. Arrows indicate nuclear bundles.

**Supplementary Figure 4.**
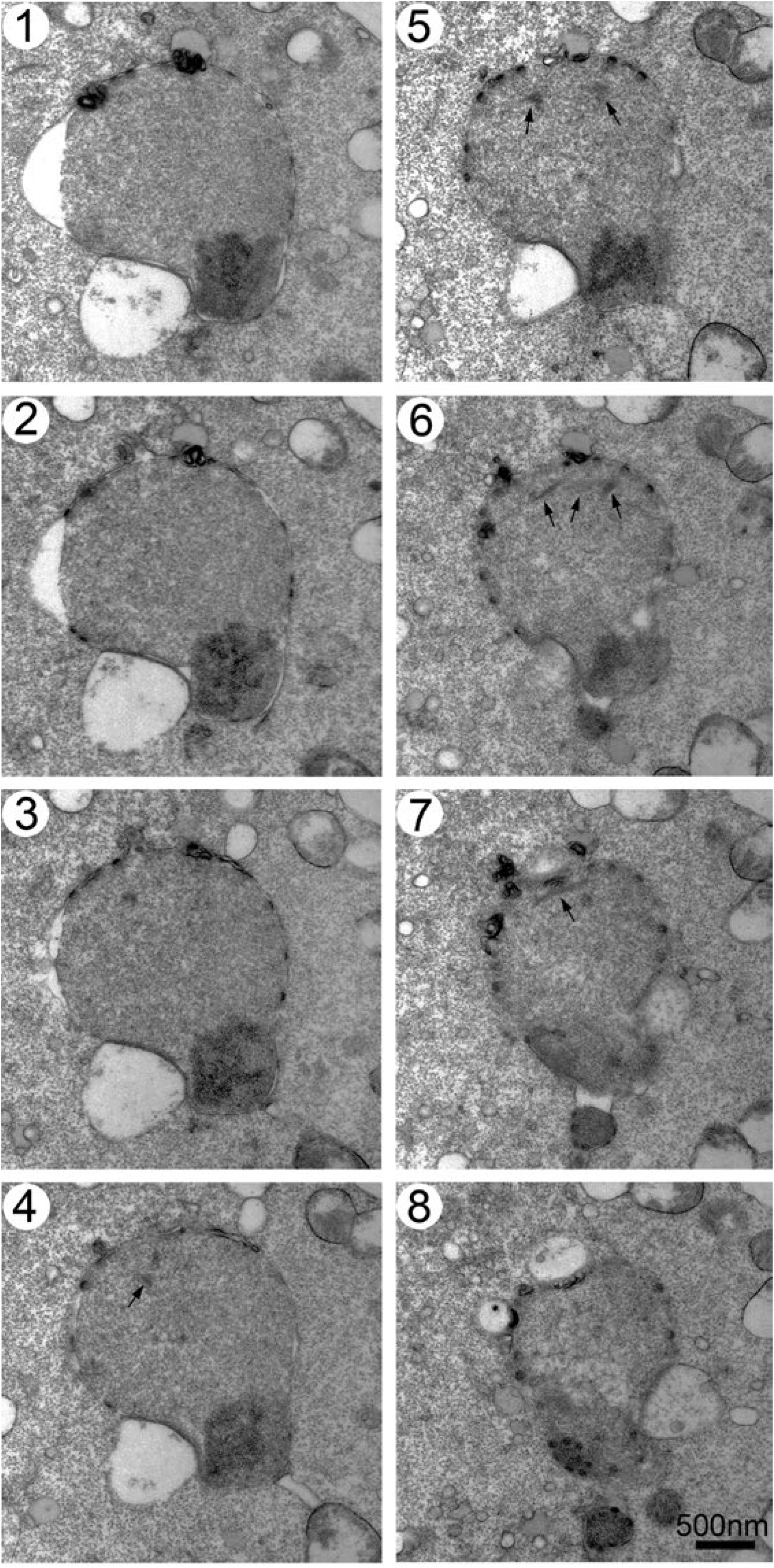
Serially sectioned EM images of a chemically fixed yeast cell in mid-prophase I. Serial sectioned TEM images of an identical cell at 4 h, cells were fixed with chemicals (glutaraldehyde and OsO_4_). Bar indicates 500 nm. Arrows indicate nuclear bundles.

**Supplementary Figure 5.**
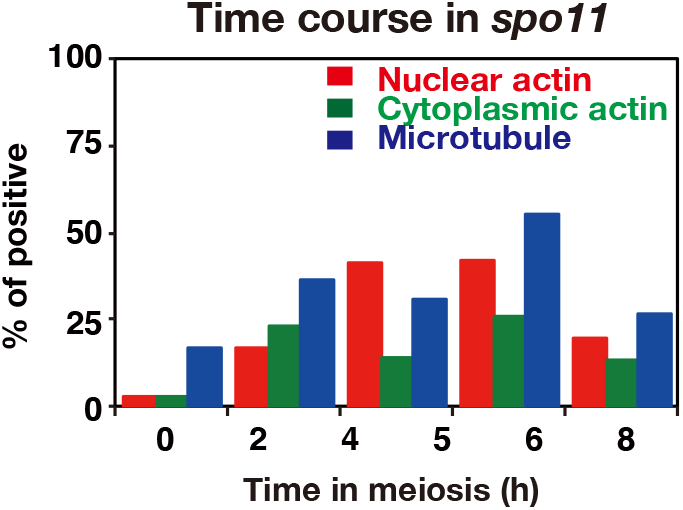
Kinetics of bundles in *spo11* cells. Kinetics of the formation of actin bundles in nuclei (red) and cytoplasm (green) as well as nuclear microtubules (blue) at different time of meiosis of the *spo11-Y135F* mutant. A number of sections containing bundles were counted and percentages of actin-positive nuclei and cytoplasm section are shown.

**Supplementary Figure 6.**
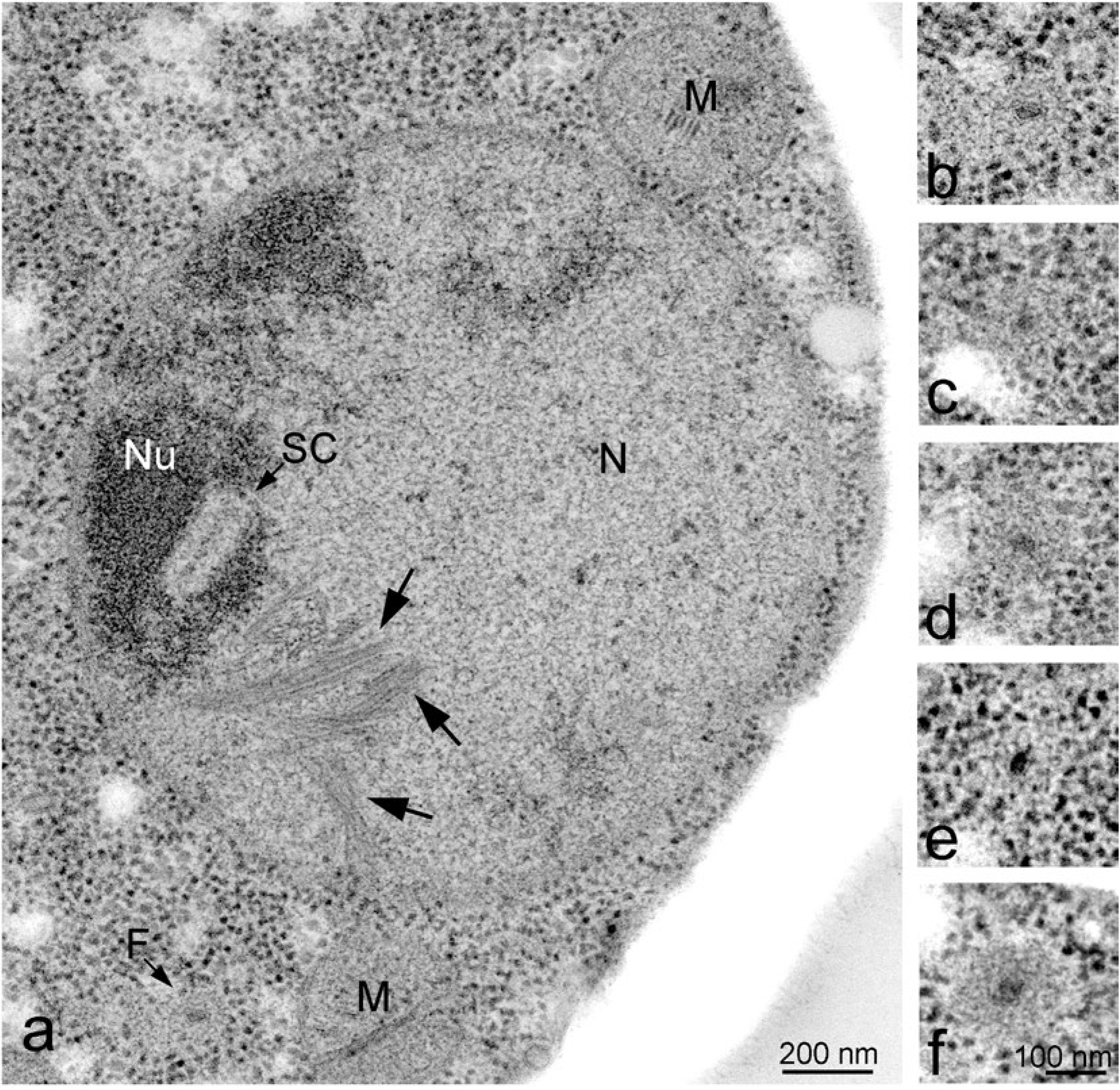
EM analysis of SC and Filasome. **a.** Representative TEM images with nuclear bundles with a possible synaptonemal complex (SC)-like structure at 5 h is shown in (a). M, mitochondrion; F, filasome; N, nucleus; Nu, nucleolus. Bar indicates 200 nm. **b-f.** Images of a filasome in a yeast diploid cell are shown. Filasomes are less dense structures with a vesicle in the center, devoid of ribosomes in cytoplasm. Bar indicates 100 nm.

**Supplementary Figure 7.**
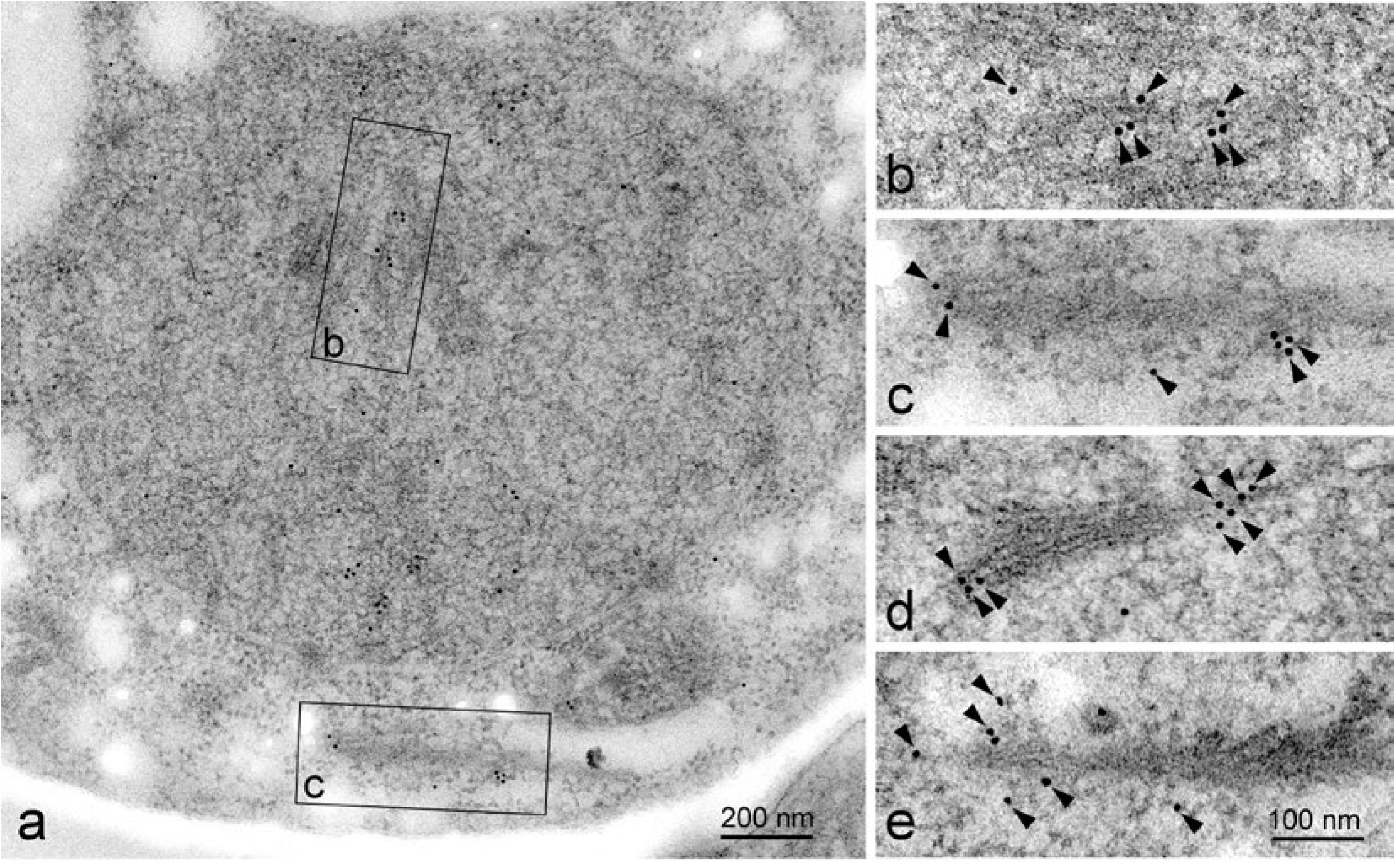
Immuno-gold labeling using anti-actin antibody. The specimens were prepared with freeze fixation. Immuno-gold labeling using anti-actin antibody was carried out as described in Materials and Methods. A representative image (a) containing actin bundles in nucleus (b) and in cytoplasm (c) in a cell at 4 h are shown with magnified images (a, b). (d, e) nuclear bundles with gold particle. The positions of gold particles are shown in arrowheads (b-e). Bar indicates 200 nm (a) and 100 nm (b-e).

